# EPF/EPFL Peptide Signaling as a Regulatory Node Linking Stomatal Development to Drought and Heat Tolerance in Tomato (*Solanum lycopersicum* L.)

**DOI:** 10.64898/2026.10.26.761844

**Authors:** Karam Mostafa, Abdullah Aydın, Ola Abd Elbar, Noor ul Ain, Musa Kavas, Hélène S Robert

**Author notes:** <u>Karam Mostafa and Abdullah Aydın</u> contributed equally to this work and shared first authorship. Corresponding authors: Musa Kavas, Hélène S Robert.

## Abstract

Stomatal development and function are tightly regulated by signaling peptides with distinct, overlapping, or antagonistic roles in gas exchange and water balance. *EPIDERMAL PATTERNING FACTOR (EPF) / EPF-LIKE* (*EPF/EPFL*) family is a founding member of the secreted peptide family that regulates stomatal patterning. The roles of *EPF/EPFL* genes in tomato responses to drought, heat, and combined drought-heat stress remain largely unexplored. To characterize the *EPF/EPFL* gene family and elucidate its potential regulatory roles in stomatal development, 15 family members were identified in *S. lycopersicum* L. All 15 candidate SlEPF proteins contained an EPF-type domain. Of these, 13 exhibited specific matches to the EPF domain, whereas SlEPF6 was classified within the EPF superfamily and SlEPF11 within the STOMAGEN-like superfamily. Phylogenetic analysis grouped *EPF* genes from diverse plant species into two major clades (A and B), each comprising distinct evolutionary subgroups. GO annotation indicated that most *SlEPF* genes were involved in stomatal complex development and guard cell differentiation, with molecular functions associated with MAP kinase activity, receptor serine/threonine kinase binding, and receptor ligand activity. Promoter analysis of *SlEPF* genes identified diverse cis-regulatory elements potentially involved in the regulation of environmental stress responses and plant development. Expression analysis of five putative stress-responsive genes (*SlEPF5, SlEPF8, SlEPF9, SlEPF10*, and *SlEPF13*) revealed distinct transcriptional patterns in the heat-tolerant drought-sensitive Money Maker (MM) and drought-tolerant Ailsa Craig (AC) tomato genotypes under drought, heat, and combined drought-heat stress. In MM, *SlEPF5* and *SlEPF13* were consistently downregulated across all stress conditions, whereas *SlEPF8* was strongly induced under drought and combined stress. In contrast, AC showed broader transcriptional activation, particularly under heat and combined stress. Overall, these contrasting expression profiles suggest genotype- and stress-dependent regulation of *SlEPF* genes, highlighting their potential involvement in tomato abiotic stress responses.

## Introduction

Global food security faces growing challenges due to rapid population growth and the intensifying impacts of climate change [1]. Under the warming climate, the increasing frequency and severity of drought and heat stress pose major threats to crop productivity, thereby exacerbating challenges to global food security [2]. A primary response of plants to drought stress is the rapid closure of stomata, which restricts carbon dioxide (CO□) diffusion into the leaves and consequently reduces the net photosynthetic rate [3, 4]. Reduced photosynthetic activity under stress conditions substantially restricts plant growth and development, as reflected by reductions in photosynthetically active leaf area, biomass accumulation, plant height, and accelerated leaf senescence [5, 6]. In addition to the direct impairment of photosynthetic processes caused by environmental stress, heat and drought stress trigger a range of physiological and cellular responses. A well-established consequence of these stresses is the excessive accumulation of reactive oxygen species (ROS), which can disrupt cellular membrane integrity and impair membrane function [7-9]. Cohen et al. (2020) demonstrated that the combined occurrence of drought and heat stress can significantly impair crop yield [10]. Consequently, there is an urgent need to develop crop varieties with enhanced resilience to this combined stress. Achieving this goal requires a comprehensive understanding of the anatomical, physiological, and molecular mechanisms underlying plant responses and adaptation to these interacting stresses [11-13].

Stomata represent a critical interface between plants and the atmosphere, that open and close to absorb the photosynthetic CO□, to regulate the exchange and movement of oxygen (O□), and to restrict water vapor during transpiration. Stomata are among the most critical anatomical features involved in plant adaptation to environmental stress. Accumulating evidence indicates that the primary physiological response to severe drought is rapid stomatal closure, which limits transpirational water loss and helps maintain plant water status [14, 15]. Stomatal closure reduces transpiration and helps regulate plant water balance; however, drought-induced stomatal closure primarily limits CO□ uptake, resulting in a reduction in photosynthetic rate [16, 17]. Mutava et al. (2015) demonstrated that drought stress significantly decreases stomatal conductance in soybean, thereby contributing to reduced photosynthetic activity [18]. In addition, drought stress can impair photosynthetic pigments and disrupt gas exchange, ultimately restricting plant growth and productivity [19]. Consequently, stomatal regulation is a key physiological process for maintaining photosynthetic capacity under stress conditions [20]. Stomatal closure is not solely dependent on drought signals originating from the leaves; rather, stomata can directly respond to soil dehydration, often before significant reductions in leaf mesophyll turgor occur [21, 22]. These findings indicate that multiple transcription factors play important roles in the regulation of stomatal development and function.

Genes regulating stomatal development and patterning have been functionally characterized [23-27]. Variation in stomatal density has been shown to affect stomatal conductance and CO_2_ assimilation rate, consequently influencing plant growth and metabolism. Stomatal patterning is precisely controlled by autocrine and paracrine signaling mediated by secreted peptides of the *EPIDERMAL PATTERNING FACTOR (EPF)* and *EPF-LIKE (EPFL)* families, which regulate stomatal development, density, and spatial distribution within the plant epidermis [28, 29]. These peptide signals are recognized by members of the *ERECTA* family of leucine-rich repeat (LRR) receptor-like kinases and/or the LRR receptor-like protein *TOO MANY MOUTHS (TMM)*, thereby modulating stomatal lineage specification, development, and spatial patterning [30, 31]. These upstream signaling events converge on a mitogen-activated protein kinase (MAPK) cascade comprising the YODA MAPKKK, the MKK4/MKK5 MAPKKs, and the MPK3/MPK6 MAPKs, which collectively regulate stomatal lineage specification, differentiation, and spatial distribution within the plant epidermis [32].

Among *EPF*/*EPF-LIKE* gene family, *STOMAGEN* uniquely functions as a positive regulator of stomatal development, whereas *EPF1* and *EPF2* act as negative regulators. STOMAGEN is structurally composed of a loop and a scaffold containing three disulfide bonds. Expressing a chimeric protein combining the *STOMAGEN* loop with the EPF2 scaffold was shown to positively regulate stomatal density in *Arabidopsis* cotyledons, suggesting that the loop confers functional specificity, while the scaffold is structurally essential for activity [33]. Several EPF transcription factors have been identified across diverse plant species, with distinct functions. *In Brassica rapa*, 14 *BrEPF* genes were identified, with potential roles in male sterility [34]. In *Zea mays* L., 16 *ZmEPF* genes [35], in *Sorghum bicolor* L., 12 *SbEPF* genes [36], in *Solanum tuberosum*, 14 *StEPF* genes [37], in *Populus trichocarpa* 15 *PtEPF* genes [38], were characterized and shown to be differentially expressed under drought conditions, suggesting their roles in drought adaptation. The involvement of EPF proteins in salt- and drought-stress response was suggested due to their differential expression pattern in *Brassica napus* L. (27 *BnEPF* genes), in oat (*Avena sativa*) (33 *AsEPF* genes) and in wheat (365 non-redundant genes) [39-41].

Although *EPF* genes play critical roles in stomatal development and plant responses to abiotic stress, their characterization in tomato (*Solanum lycopersicum* L.) remains limited. In this study, we aim to investigate the *EPF* gene family in tomato and elucidate its role in stomatal development and regulation under drought, heat, and combined drought-heat stress conditions using drought-tolerant and drought-sensitive tomato varieties.

## Material and Methods

### Identification and Retrieval of the *EPF/EPFL* Gene Family in Tomato

A genome-wide approach was employed to identify the *EPF* gene family in tomato. Candidate SlEPF proteins were identified using The Hidden Markov Model (HMM-) and BLASTP-based searches against the tomato proteome, using the conserved EPF domain and previously characterized EPF proteins as queries [42]. Additionally, keyword searches were performed against selected *Solanum lycopersicum* genome in the Phytozome v14 database (https://phytozome-next.jgi.doe.gov/) to identify potential genes missed by sequence-based searches. All candidate protein sequences were subsequently examined to confirm the presence of the characteristic EPF domain using InterPro (https://www.ebi.ac.uk/interpro/) and Pfam (http://pfam-legacy.xfam.org/) databases [43, 44]. Redundant sequences and non-functional protein models were excluded from further analysis. The confirmed *SlEPF* genes were retrieved together with their corresponding coding and genomic sequences from the Phytozome v14 database. The identified genes were systematically named according to their chromosomal positions and genomic organization. The physicochemical properties of the predicted SlEPF proteins, including amino acid length, molecular weight, and theoretical isoelectric point (pI), aliphatic index, grand average hydropathy (GRAVY) using the ExPASy ProtParam tool (http://www.expasy.ch/tools/pi_tool.html). Subcellular localization and other relevant protein characteristics were subsequently predicted using the Plant-mPLoc server (http://www.csbio.sjtu.edu.cn/bioinf/plant-multi/) [45]. Signal peptides and their corresponding cleavage sites were predicted from the full-length amino acid sequences of SlEPF proteins using SignalP v6.0 (https://services.healthtech.dtu.dk/services/SignalP-6.0/) [46]. The analysis was performed using the eukarya organism group and the slow-sequential prediction mode. SignalP outputs were converted into a binary classification for tabulation: proteins assigned to the Signal Peptide (Sec/SPI) class with an SP probability greater than 0.50 were recorded as “Yes” whereas proteins assigned to the other class were recorded as “No” for proteins classified as Sec/SPI, the predicted signal peptide region and cleavage-site position were recorded according to the SignalP output.

### Structural Features and Phylogenetic Classification of the *SlEPF* Gene Family

The structural organization of the *SlEPF* genes was characterized to elucidate patterns of structural diversification within the *SlEPF* gene family and to provide insights into its potential evolutionary history. Conserved motifs were identified from the SlEPF protein sequences using the Multiple Em for Motif Elicitation (MEME) Suite (https://meme-suite.org/meme/tools/meme) [47]. The MEME analysis was performed by specifying a maximum of 15 conserved motifs, with each protein sequence allowed to contain no more than one occurrence of each motif. Motif lengths were restricted to 6–50 amino acids, whereas all other parameters were retained at their default settings. To ensure the robustness of motif prediction, the identified motifs were individually evaluated, and only those exhibiting an E-value < 10□¹□ were considered significant and retained for downstream analyses of the SlEPF proteins. Conserved domains within the SlEPF protein sequences were identified using the NCBI Conserved Domain Database (CDD) through the CD-Search and Batch CD-Search tools (https://www.ncbi.nlm.nih.gov/Structure/cdd/wrpsb.cgi) [48]. The detected domains were subsequently examined and compared among the SlEPF proteins to assess their structural conservation and potential functional relationships. Additionally, the exon–intron architecture of the *SlEPF* genes was determined based on the corresponding GFF3 genome annotation file. The resulting motif, conserved domains and gene structures were subsequently visualized using TBtools [49].

To investigate the evolutionary relationships and divergence of the *EPF* gene family across plant species, putative EPF protein sequences from *Arabidopsis thaliana, Nicotiana benthamiana, Phaseolus vulgaris, Triticum aestivum, Musa acuminata, Gossypium hirsutum, Solanum tuberosum, Fragaria vesca, Vitis vinifera, Zea mays, Populus trichocarpa, Vaccinium darrowii, and Oryza sativa* were retrieved from the Phytozome v14 database. The resulting dataset was used to reconstruct the evolutionary relationships of EPF proteins within and among species and to provide a phylogenetic framework for subsequent comparative and synteny analyses. Multiple sequence alignment of the EPF protein sequences was performed using ClustalW 2.0 implemented in MEGA 12 [50]. Phylogenetic relationships were reconstructed using the neighbor-joining (NJ) method, and the reliability of the inferred topology was evaluated using 1,000 bootstrap replicates. The resulting phylogenetic tree was subsequently visualized and graphically refined in R using the ggtree package.

### Three-dimensional Structure Visualization and Prediction of Putative Disulfide Bonds

To gain deeper insight into the domain organization, three-dimensional protein structure, and structural stability of the SlEPF proteins, their predicted three-dimensional structures and putative disulfide bonds were analyzed. Putative three-dimensional structures of the 15 SlEPF proteins were predicted using AlphaFold2 [51]. All parameters were kept at their default settings, except num_relax, which was set to 1. The resulting relaxed models were obtained in PDB format and visualized using UCSF ChimeraX [52]. Proteins were displayed using a cartoon representation, while the annotated domain regions were distinguished using a consistent color scheme. Cysteine residues were displayed as sticks and individually labelled to facilitate the examination of their spatial distribution. Putative intramolecular disulfide connectivity was evaluated by measuring the distances between the side-chain sulfur atoms (Sγ) of spatially adjacent cysteine residues using the distance measurement function in ChimeraX.

### Protein-Protein Interaction and Gene Ontology Analysis

Protein-protein interaction (PPI) analysis of the SlEPF family was performed using the Search Tool of the Retrieval of Interacting Genes (STRING, V 12.5) web portal (http://string-db.org) [53]. All SlEPF proteins were submitted simultaneously to generate a combined interaction network in *S. lycopersicum*. The network was constructed using the Full STRING network option, with network edges displayed according to interaction confidence. A minimum required interaction score of 0.7 was applied to retain only relatively well-supported associations. To expand the initial network and incorporate additional functionally associated proteins, the “More” option was applied twice. All remaining STRING parameters were kept at their default settings. Functional enrichment analysis was subsequently performed directly on the resulting expanded interaction network. Gene Ontology (GO) enrichment was evaluated separately for the three major GO categories: Biological Process (BP), Molecular Function (MF), and Cellular Component (CC). Significantly enriched GO terms identified by STRING were used to characterize the major biological processes, molecular functions, and subcellular associations represented within the SlEPF interaction network.

### Analysis of Cis-Regulatory Elements in the Promoter Regions of *SlEPF* genes family

Promoter regions corresponding to the 2,000-bp sequences upstream of each *SlEPF* gene were retrieved from the Phytozome V14 and used for cis-regulatory analysis. Putative cis-acting regulatory elements were identified using the PlantCARE database (https://bioinformatics.psb.ugent.be/webtools/plantcare/html/) [54]. The identified elements were subsequently classified according to their reported functional associations into major categories, including light responsiveness, phytohormone responsiveness, stress responsiveness, development/metabolism, and circadian regulation. For each *SlEPF* promoter, both the total occurrence of individual cis-elements and the number of distinct cis-regulatory element types were determined. To further characterize the transcriptional regulatory landscape of the *SlEPF* family, putative transcription factor-binding sites (TFBSs) within the same promoter regions were predicted using PlantPAN 4.0 (https://plantpan.itps.ncku.edu.tw/plantpan4/) [55]. Predicted binding motifs were grouped according to their corresponding transcription factor families, and the number of motif occurrences was calculated for each promoter. In addition, the total number of motif-position associations were determined to compare the overall density of predicted TFBSs among *SlEPF* promoters. The resulting cis element and TFBS distributions were summarized and visualized as heatmaps together with promoter and motif level abundance plots.

### Tissue-specific expression analysis

Tissue-specific expression profiles of the 15 identified *SlEPF* genes were examined using the Tomato eFP Browser provided by the Bio-Analytic Resource for Plant Biology (BAR), University of Toronto (https://bar.utoronto.ca/) [56]. Each gene was queried using its corresponding Solyc identifier, and publicly available expression values across tomato organs and developmental stages were retrieved from the tomato atlas dataset [57]. The values were compiled into a gene-by-sample matrix and visualized as a heatmap to compare expression patterns among family members. *SlEPF9* was excluded from the heatmap because expression data for this gene were unavailable in the selected dataset.

### Plant Materials and Experimental Growth Conditions

Two well-characterized commercial tomato cultivars were selected as the experimental plant materials heat-tolerant drought-sensitive Money Maker (MM) [58], and drought-tolerant Ailsa Craig (AC) tomato genotypes [59]. Seeds were sown individually in polypropylene pots (11 × 11 × 12 cm) containing a 2:1 (v/v) mixture of peat moss and vermiculite. Four biological replicates were maintained for each treatment. All plants were cultivated under controlled environmental conditions in a walk-in Fytoscope growth chamber (FS-WI; Photon Systems Instruments [PSI], Czech Republic). Plants were grown under a 16 h light/8 h dark photoperiod using LED illumination at a photosynthetic photon flux density (PPFD) of 150 µmol m□² s□¹. Relative humidity was maintained at 35–45%, with air temperatures of 26 ± 2 °C during the light period and 18 ± 2 °C during the dark period. These controlled conditions were maintained throughout plant growth.

At the two- to three-leaf stage, tomato plants were randomly assigned to four treatments, with four biological replicates per treatment: (i) Control, plants maintained under standard conditions and irrigated daily with 150 mL water; (ii) heat stress, plants exposed to 36 °C for 6 h day□¹ while receiving 150 mL water daily; (iii) drought stress, plants irrigated with 30 mL water pot□¹ day□¹ to maintain soil moisture at approximately 20% of field capacity; and (iv) combined heat and drought stress, plants subjected simultaneously to 36 °C for 6 h day□¹ and irrigation with 30 mL water pot□¹ day□¹, maintaining soil moisture at approximately 20% of field capacity.

### RNA Extraction and Expression Profiling of Putative Abiotic Stress-Responsive *EPF* Genes

At the molecular level, five putative abiotic stress-responsive *EPF* genes (*SlEPF5*, *SlEPF8*, *SlEPF9*, *SlEPF10*, and *SlEPF13*) were selected for expression analysis in tomato under the different abiotic stress conditions. The genes were selected based on the presence and abundance of stress-responsive cis-regulatory elements (CERs) in their promoter regions and PPI analysis. The sequences of all primers used in this study are provided in Supplementary Tables S1. Leaf samples were collected from each tomato genotype under control and abiotic stress conditions, immediately flash-frozen in liquid nitrogen, and stored at −80 °C until further processing. Total RNA was extracted using the NucleoSpin™ RNA Plant kit (Macherey-Nagel, Germany) according to the manufacturer’s instructions. RNA concentration and purity were determined spectrophotometrically using a NanoDrop™ 2000/2000c spectrophotometer (Thermo Fisher Scientific, USA), while RNA integrity was assessed by electrophoresis on a 1.2% (w/v) agarose gel. First-strand complementary DNA (cDNA) was synthesized from 1,000 ng of total RNA using Moloney murine leukemia virus reverse transcriptase (M-MLV RT), following the manufacturer’s protocol. Quantitative PCR (qPCR) was performed using LightCycler® 480 SYBR Green I Master Mix (Roche, Switzerland), with 1 µL of cDNA in a final reaction volume of 20 µL. Amplification was carried out using a LightCycler 96 machine (Roche, Switzerland). The thermal cycling program consisted of an initial preincubation step at 95 °C for 10 min, followed by 40 amplification cycles comprising denaturation at 95 °C for 10 s, annealing at 60 °C for 10 s, and extension at 72 °C for 20 s. Relative transcript abundance was calculated using the 2^−^ΔΔCt^ method. Gene expression levels were normalized against two reference genes: tomato ACTIN2 (*SlACT2; Solyc11g005330*) and ELONGATION FACTOR 1-ALPHA (*EF1*α*; Solyc06g009970*) (Supplementary Tables S1). These reference genes were used to account for variation in cDNA input and other technical factors during expression analysis.

## Results

### Systematic Identification and Physicochemical Characterization of *SlEPF* Genes

A comprehensive BLAST-P search against the predicted *Solanum lycopersicum* proteome available in the Phytozome v14 database identified 15 genes encoding putative SlEPF proteins (Supplementary File 1). The candidate sequences were further examined for the presence of the characteristic conserved EPF to confirm their identity as SlEPF family members. Sequences lacking the conserved EPF domain were excluded from subsequent analyses. The complete set of identified *SlEPF* genes, together with their corresponding annotation information, is presented in Supplementary Table S2. The identified *SlEPF* members were systematically assigned numerical identifiers (*SlEPF1* to *SlEPF15*) based on their chromosomal locations, following their sequential physical order along the chromosomes of *S. lycopersicum* (Supplementary File 2, Figure S1). This chromosome-based nomenclature was adopted to provide a consistent and systematic designation for the members of the *SlEPF* gene family. The predicted SlEPF proteins exhibited considerable variation in sequence length, ranging from 55 amino acids in SlEPF13 to 177 amino acids in SlEPF2, with an average length of approximately 107 amino acids. This pronounced variation in protein size suggests a structural diversity among the members of the SlEPF family in tomato. Analysis of the predicted physicochemical properties of the SlEPF proteins revealed marked variation among family members. The estimated molecular weight ranged from 5.95 kDa for SlEPF13 to 39.91 kDa for SlEPF4, with an average of approximately 11.77 kDa. Similarly, the predicted isoelectric point (pI) values ranged from 8.70 in SlEPF3 to 10.22 in SlEPF8, with a mean value of approximately 9.93. Overall, the variation in molecular weight and pI values indicates physicochemical diversity among the SlEPF proteins. The GRAVY analysis revealed predominantly negative values across the SlEPF protein family, with the exceptions of SlEPF10, SlEPF11, SlEPF13, and SlEPF15, which exhibited positive GRAVY scores. The negative GRAVY values observed for most SlEPF proteins indicate a predominantly hydrophilic nature, whereas the positive values detected in the aforementioned proteins suggest comparatively greater hydrophobicity (Supplementary Table S2). *In silico* prediction of subcellular localization places 14 of the 15 SlEPF proteins to the nucleus, either exclusively (SlEPF1-6 and SlEPF14) or in combination with other compartments. SlEPF9 was uniquely predicted to be chloroplast localized. Additional predicted sites included the chloroplast (SlEPF7-12), cell membrane (SlEPF10, 12, 13, 15), mitochondrion (SlEPF11), Golgi apparatus (SlEPF13), and cell wall (SlEPF15). The diversity of predicted localizations, particularly among SlEPF10/11/12/13/15, suggests possible functional divergence within the protein family. SignalP 6.0 classified nine SlEPF proteins as containing an N-terminal Sec/SPI signal peptide, seven of which exhibited high-confidence predictions. SlEPF10 and SlEPF15 showed comparatively lower-confidence signal peptide predictions, with the predicted cleavage site of SlEPF15 being particularly atypical. Thus, the predicted extracellular localization of several SlEPF members was not accompanied by a detectable canonical N-terminal signal peptide (Supplementary Table S2).

### Conserved Structural Characteristics and Evolutionary Relationships of the *SlEPF* Gene Family

To characterize the *SlEPF* gene family at both the nucleotide and protein levels, the 15 identified *SlEPF* members were systematically analyzed based on their phylogenetic relationships, sequence motifs, conserved domains and exon–intron structures (Figure 1). The phylogenetic analysis classified the genes into six distinct subgroups (Figure 1A). Conserved motifs in the SlEPF proteins were identified with MEME (v5.5.9; zero-or-one occurrence per sequence, motif width 6-50 aa, maximum of 15 motifs). Fifteen motifs were returned (Figure 1B and Supplementary Table S3), but only two were statistically significant. Motif 1 (28 aa; E = 4.4 × 10□□¹) was found in 12 of the 13 analyzed sequences. It was also detected in the remaining sequence in the subsequent scan (p < 10□□). Motif 2 (21 aa; E = 4.0 × 10□□□) was present in all 13. Together, these motifs formed the conserved cysteine-rich core of the family. Motif 1 contains four invariant cysteines in a C-x□-C-x□-C-x□-C arrangement, flanked by conserved Gly, Ser, and Pro residues. Motif 2 lies at the C-terminus and contains a conserved C-x-C module and a conserved aromatic residue (Trp/Tyr). In every sequence, Motif 1 preceded Motif 2. The two motifs together contribute six conserved cysteines, consistent with the cysteine-rich domain characteristic of EPF peptides. The remaining motifs (Motifs 3-15) were short, were found in only two to four sequences, and had weak statistical support (E = 1.9 to 8.2 × 10^-^³). The conservation of the C-terminal cysteine-rich region, together with divergent N-terminal and intervening regions, suggests that the SlEPF family has diversified while retaining its core structure. To confirm that the candidate genes encode EPF proteins, the 15 predicted tomato protein sequences were screened for conserved domains with the NCBI Batch CD-Search tool against the Conserved Domain Database. Conserved domain analysis showed that all 15 candidate SlEPF proteins contain an EPF-type domain (Figure 1C). Thirteen proteins carried a specific hit to the EPF domain (pfam17181; cl28844), with E-values from 9.35 × 10□¹□ to 7.26 × 10□□ and bitscores from 45.5 to 72.5. SlEPF6 matched the EPF superfamily (cl28844; E = 2.50 × 10□□), and SlEPF11 matched the STOMAGEN-like superfamily (cl25585; E = 1.51 × 10□³□), consistent with its relationship to STOMAGEN/EPFL9. The EPF domain was 47-78 amino acids long and was located in the central region of most proteins. SlEPF3 showed a marginal hit to the 7tm_GPCRs superfamily (E = 3.2 × 10□³), which was disregarded. SlEPF4 also contained an mTERF domain (E = 9.24 × 10□¹□) in addition to its C-terminal EPF domain (residues 301-347), suggesting an atypical domain architecture. Overall, these results support the classification of the candidates as members of the SlEPF family. Gene structure analysis showed that *SlEPF9* and *SlEPF3* are intronless, *SlEPF1*, *SlEPF10* and *SlEPF11* contain a single intron, and the remaining genes are multi-exonic, with genomic lengths ranging from about 0.15 to 6.8 kb (Figure 1D). *SlEPF4* and *SlEPF8* contain long introns. The exon–intron organization was generally similar within clades. Overall, the conserved motif and domain architectures observed among closely related family members were generally consistent with their phylogenetic relationships, while the distinctive structures of a small number of proteins indicate structural diversification within the SlEPF family.

**Figure 1.**
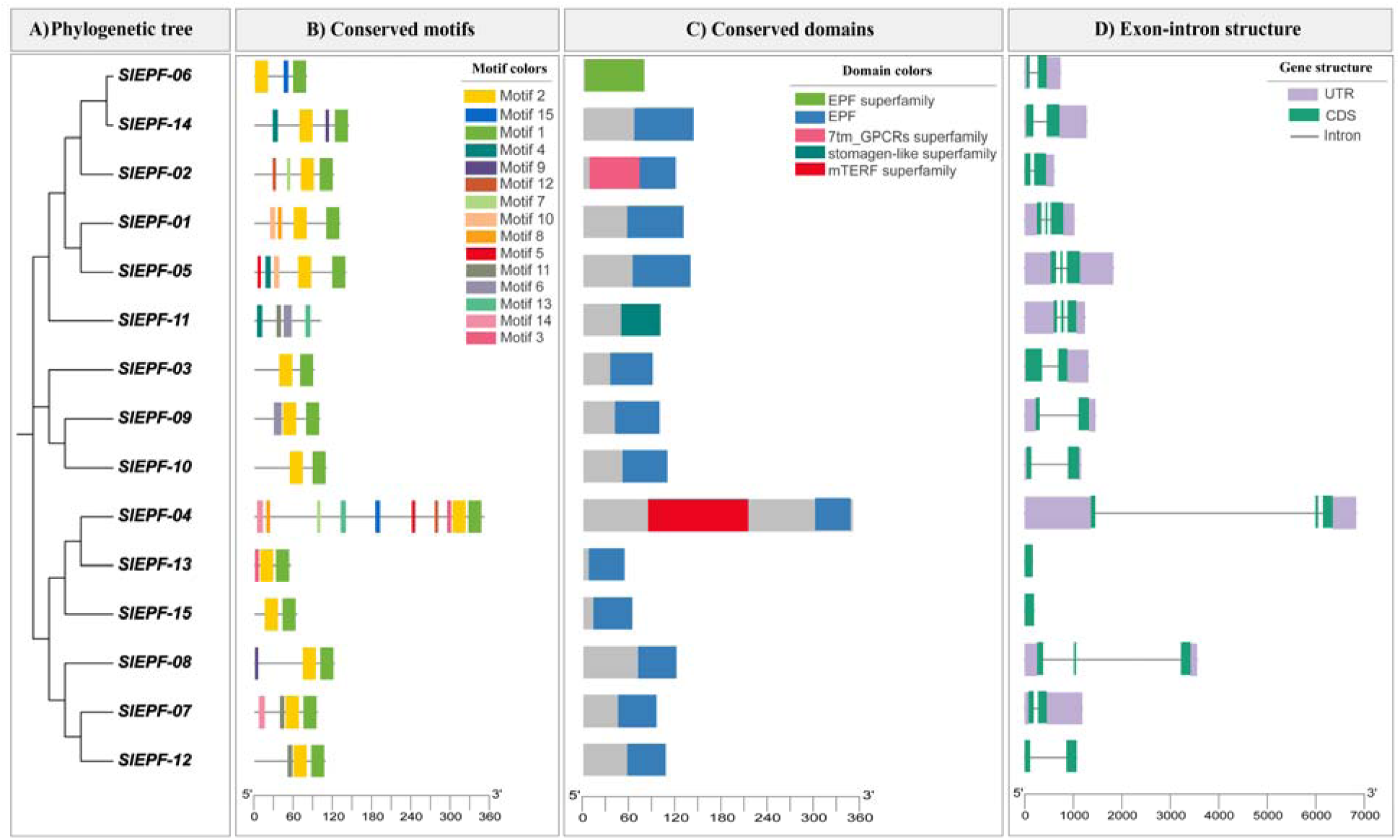
Phylogenetic relationships, conserved motifs, conserved domains and gene structures of the 15 *SlEPF* family members. A) Phylogenetic tree of the SlEPF proteins and (B) distribution of the 15 conserved motifs identified by MEME. Motifs 1-15 are shown as colored boxes, and the scale shows protein length in amino acids (aa). C) Conserved domains identified by NCBI CD-Search. The EPF domain is shown in blue, the EPF superfamily in green, the STOMAGEN-like superfamily in teal, the 7tm_GPCRs superfamily in pink and the mTERF superfamily in red. Gray indicates the remaining protein sequence. D) Exon-intron structure of the *SlEPF* genes. Purple boxes indicate UTRs, green boxes indicate CDS (exons), and gray lines indicate introns. The scale shows gene length in base pairs (bp). The composite figure was visualized using TBtools.

To investigate the evolutionary relationships of the tomato *EPF* family, a phylogenetic tree was built from 128 EPF protein sequences from 14 plant species, *Gossypium hirsutum* (21), *Triticum aestivum* (16), *Populus trichocarpa* (10), *Zea mays* (10), *Phaseolus vulgaris* (9), *Musa acuminata* (8), *Arabidopsis thaliana* (7), *Vitis vinifera* (7), *Solanum tuberosum* (6), *Oryza sativa* (6), *Fragaria vesca* (6), *Nicotiana benthamiana* (4), *Vaccinium darrowii* (3) and 15 from tomato (Figure 2). The tree separated into two large clades (Clade A, 81 proteins; Clade B, 46 proteins) and the long-branched SlEPF11. The 15 SlEPF proteins were distributed across both clades. SlEPF7, SlEPF8 and SlEPF12 grouped with AtEPF1/AtEPF2 or AtEPF4/AtEPF7 and with potato homologs (support 0.98-1.0), whereas SlEPF15 grouped with StEPF3 and the AtEPF3-containing clade, and SlEPF4 and SlEPF13 grouped with the AtEPF6-containing clade. SlEPF1/5, SlEPF6/14, SlEPF4/13 and SlEPF9/10 formed tomato-specific sister pairs, and SlEPF2, SlEPF9 and SlEPF10 formed a tomato-specific group without orthologs in the other species, suggesting lineage-specific duplication. Monocot proteins (maize, rice and wheat) mostly clustered separately from eudicot proteins, indicating family diversification after the monocot-eudicot split. Many deep nodes had low support, so only the terminal relationships were interpreted. Additionally, we have noticed that the SlEPF6/SlEPF14, SlEPF1/SlEPF5, SlEPF10/SlEPF9, SlEPF4/SlEPF13, and SlEPF7/SlEPF12 pairs, which belong to the same phylogenetic clades in Figure 2, exhibit similar gene structures and conserved motif distributions, as illustrated in Figure 1. Although SlEPF7 and SlEPF12 also shared similar structural features, they occupied distinct positions within the phylogenetic tree, indicating a divergence in their evolutionary relationships (Figure 2).

**Figure 2.**
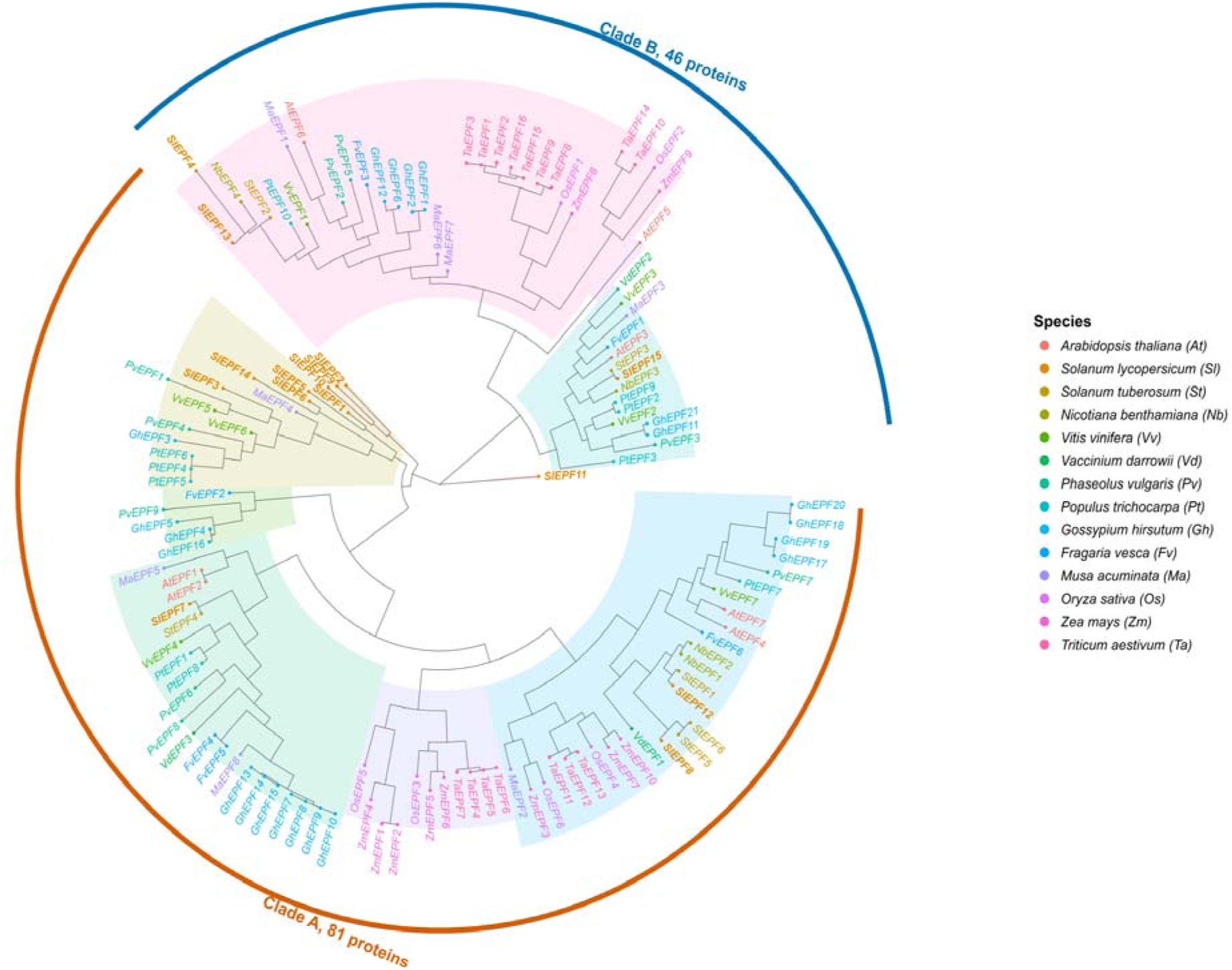
Phylogenetic relationships of the EPF family in tomato and other plant species. The tree was constructed from 128 EPF protein sequences using the maximum likelihood method in MEGA12. The proteins were divided into two major clades: Clade A (orange line), comprising 81 proteins, and Clade B (blue highlight), comprising 46 proteins. Tip colors indicate species, and the shaded sectors indicate major clades. Tomato members (SlEPF1-15) are shown in bold italics. Species abbreviations: At, *Arabidopsis thaliana*; Sl, *Solanum lycopersicum*; St, *Solanum tuberosum*; Nb, *Nicotiana benthamiana*; Vv, *Vitis vinifera*; Vd, *Vaccinium darrowii*; Pv, *Phaseolus vulgaris*; Pt, *Populus trichocarpa*; Gh, *Gossypium hirsutum*; Fv, *Fragaria vesca*; Ma, *Musa acuminata*; Os, *Oryza sativa*; Zm, *Zea mays*; Ta, *Triticum aestivum*. The figure was generated with ggtree in R.

### Predicted 3D Structures and Conserved Cysteine Architecture of SlEPF Proteins

Multiple sequence comparison revealed pronounced conservation within the C-terminal regions of EPF proteins. In tomato, 13 of the 15 SlEPF proteins terminated with a proline residue. A similar tendency was observed among *Arabidopsis* homologs, although the terminal proline was not universal. Notably, several phylogenetically related members displayed highly conserved terminal motifs, including YPVP, F/YMP, YNP and HVPS-containing sequences, suggesting that C-terminal conservation extends beyond a single terminal residue and is associated with specific EPF subgroups. The predicted three-dimensional structures were subsequently examined to assess the spatial organization of conserved cysteine residues. Putative intramolecular disulfide bonds were assigned when the distances between the sulfur atoms (Sγ) of paired cysteine residues were compatible with disulfide-bond geometry (Figure 3). The measured Sγ-Sγ distances were consistently close to approximately 2.0 Å, supporting the structural plausibility of the predicted cysteine pairings (Supplementary Table S4). Most SlEPF proteins contained three putative disulfide bonds, whereas two and four putative bonds were identified in SlEPF5 and SlEPF15, respectively. Nearly all predicted disulfide bonds were located within the annotated EPF-related domain regions. The only exception was the Cys39-Cys48 pair in SlEPF15, which was positioned outside the annotated EPF domain. Collectively, these findings reveal a largely conserved cysteine-connectivity pattern among SlEPF proteins while also indicating member-specific structural variation in SlEPF5 and SlEPF15.

**Figure 3.**
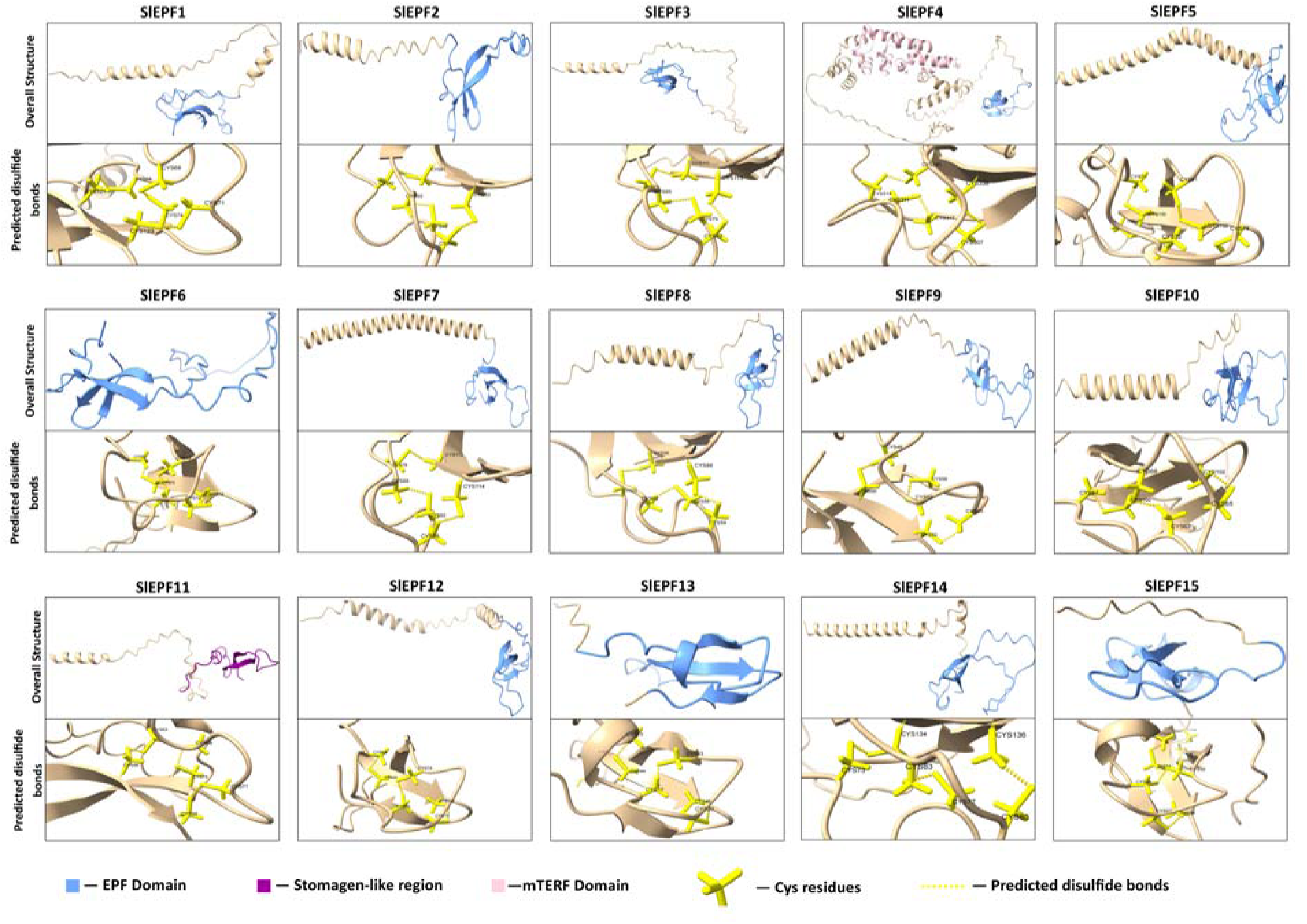
Predicted three-dimensional structures and disulfide-bond architecture of SlEPF proteins. Upper panels show the overall structures of SlEPF1-SlEPF15 predicted using AlphaFold2 and subjected to structural relaxation. EPF domains are highlighted in blue, the STOMAGEN-like region in purple, and the mTERF domain in pink, while the remaining regions are shown in beige. Lower panels provide enlarged views of the cysteine-rich regions. Cysteine residues are displayed as yellow sticks, and yellow dashed lines indicate predicted intramolecular disulfide bonds based on S-S distances compatible with disulfide-bond formation.

### Predicted protein-protein interaction network and functional annotation of SlEPF proteins

PPI analysis using the STRING database was accompanied by Gene Ontology (GO) enrichment analysis to characterize the biological processes, molecular functions, and cellular components associated with SlEPF proteins. PPI analysis using STRING predicted a network of 24 nodes, comprising 15 SlEPF proteins and nine additional interacting partners. Two SlEPF proteins, SlEPF11 and SlEPF5, showed an interaction with each other in the predicted PPI network. Only two SlEPFs (SlEPF15 and SlEPF13) were connected to other proteins; the remaining eleven were isolated nodes with no predicted interactions.

SlEPF13 and SlEPF15 were linked to a densely connected group of partners (A0A3Q7G285 (Serine/Threonine-protein), A0A3Q7HPD8 (Serine/Threonine-protein), A0A3Q7J9L5 (LRR receptor-like serine/threonine-protein kinase ERECTA)) and, in addition, for SlEPF15, A0A3Q7G821 (Epidermal patterning factor-like protein). A second cluster of three interconnected proteins (A0A3Q7IQ87, A0A3Q7G3I6 and A0A3Q7GQI0 (mitogen-activated protein kinase kinase kinase)) was also connected to this hub through their interaction with Serine/Threonine proteins (A0A3Q7G285, A0A3Q7HPD8). The links with A0A3Q7G285 and A0A3Q7HPD8 forms the central hub of the network, to which the proteins A0A3Q7GNR8 and A0A3Q7GBA2 (Uncharacterized proteins) were connected through their interaction with A0A3Q7G285. SlEPF11 interacted only with SlEPF5, through a low-confidence edge (Figure 4).

**Figure 4.**
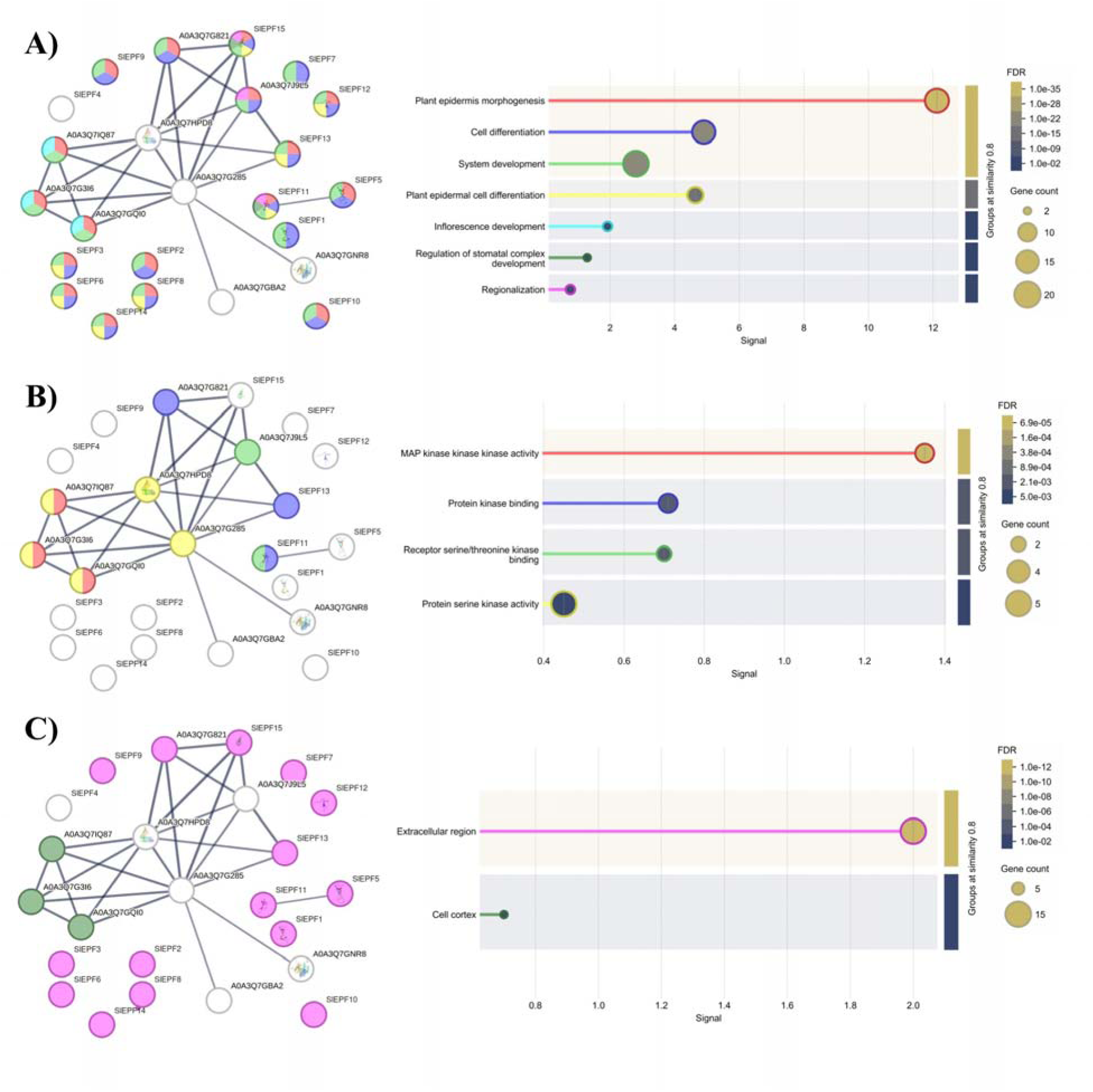
PPI network and GO enrichment analysis of the SlEPF protein family. PPI networks were constructed using the STRING database, and functional enrichment was evaluated according to the three major GO categories: (A) Biological Process, (B) Molecular Function, and (C) Cellular Component. In each panel, the STRING-derived interaction network is shown on the left, whereas the corresponding significantly enriched GO terms are presented on the right. Colored sectors within network nodes indicate the GO terms associated with individual proteins, while uncolored nodes indicate proteins not assigned to the displayed enriched terms. In the enrichment plots, the x-axis represents the enrichment signal, circle size corresponds to the number of genes associated with each GO term, and circle color represents the false discovery rate (FDR). GO terms were grouped based on a similarity threshold of 0.8. Biological Process enrichment was dominated by developmental categories, including plant epidermis morphogenesis, cell differentiation, and system development, whereas Molecular Function enrichment primarily involved kinase-related binding activities. Cellular Component analysis predominantly associated SlEPF proteins with the extracellular region.

The Biological Process analysis revealed that most SlEPF family members were associated with developmental processes, particularly plant epidermis morphogenesis, cell differentiation, and system development (Figure 4A). Among these terms, plant epidermis morphogenesis represented the most strongly enriched biological process. Notably, SlEPF4, which represents a divergent member of the SlEPF family, was not associated with any of the significantly enriched biological processes identified in this analysis. In contrast, SlEPF11 and SlEPF15 were additionally associated with the regulation of stomatal complex development, distinguishing them from the other SlEPF family members and suggesting a potentially more specific involvement in stomatal development. For the Molecular Function category, most SlEPF proteins were not associated with any significantly enriched functional terms. However, SlEPF11 and SlEPF13 were associated with protein kinase binding activity, while SlEPF11 was additionally linked to receptor Serine/Threonine kinase binding. These associations indicate that SlEPF11, and to a lesser extent SlEPF13, may participate in kinase-mediated signaling interactions that are not broadly shared across the SlEPF family (Figure 4B). For the Cellular Component category, all SlEPF family members, with the exception of the divergent SlEPF4 protein, were associated with the extracellular region. This pattern is consistent with the predominantly extracellular localization expected for EPF family proteins and further highlights SlEPF4 as an atypical member of the family (Figure 4C).

### Cis-regulatory elements and transcription factor-binding site profiles of *SlEPF* promoters

Analysis of the upstream promoter regions of *SlEPF* genes revealed a diverse repertoire of cis-regulatory elements associated with environmental and developmental responses (Figure 5A). PlantCARE analysis showed that light-responsive elements were the most abundant category, with a total of 52 occurrences across the *SlEPF* promoters, followed by ABA-responsive elements with 30 occurrences. Several additional phytohormone-responsive elements, as well as cis-elements associated with stress responses, development/metabolism, and circadian regulation, were also identified. Among the stress-related elements, those associated with anaerobic induction were particularly abundant, indicating a substantial representation of stress-responsive regulatory motifs within the promoters. The composition and abundance of these elements varied considerably among individual *SlEPF* genes, with the *SlEPF14* promoter displaying the greatest diversity of non-redundant cis-element annotations. Overall, these results indicate substantial heterogeneity in the putative regulatory architecture of *SlEPF* promoters.

**Figure 5.**
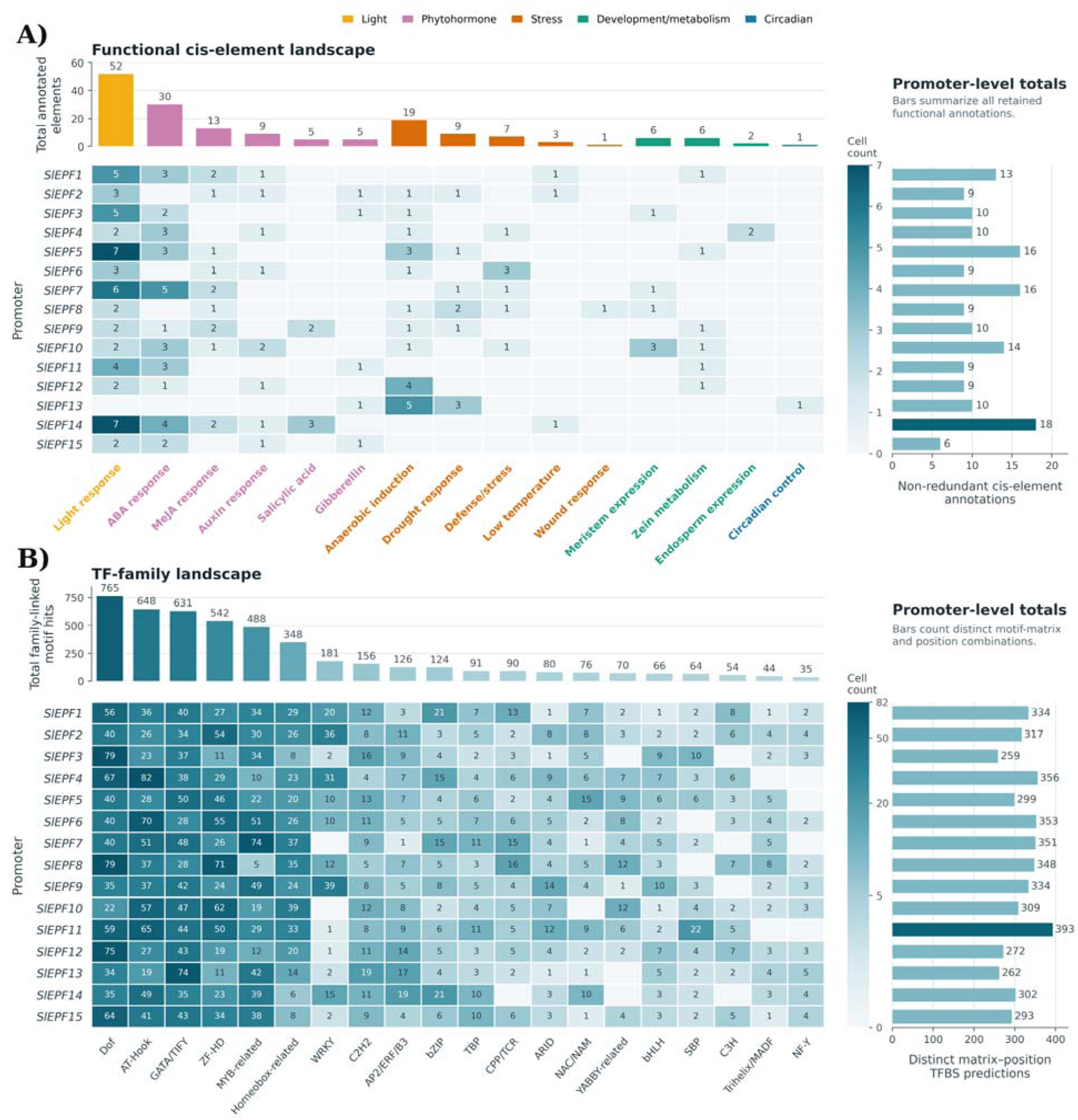
Cis-regulatory element and TFDBS in the promoter regions of *SlEPF* genes. (A) Distribution of cis-regulatory elements identified in the approximately 2-kb upstream promoter regions of *SlEPF* genes using PlantCARE. Identified elements were grouped according to their putative functional categories, including light responsiveness, phytohormone responsiveness, stress responsiveness, development/metabolism, and circadian regulation. The heatmap shows the number of individual cis elements detected in each promoter, whereas the vertical bars above indicate their overall abundance across the *SlEPF* family. Horizontal bars on the right represent the total number of non-redundant cis-element annotations detected for each promoter. (B) Distribution of predicted TFBSs in *SlEPF* promoters based on PlantPAN 4.0 analysis. The heatmap illustrates the number of predicted binding sites belonging to different transcription factor families within each promoter, while the vertical bars above indicate the total abundance of each TFBS category across the analyzed promoters. Horizontal bars on the right show the total number of distinct motif-position TFBS predictions for each *SlEPF* promoter. Color intensity in both heatmaps reflects the number of predicted cis-elements or TFBSs.

Consistent with the diversity observed in cis-regulatory elements, PlantPAN-based analysis identified numerous putative transcription factor-binding sites across all *SlEPF* promoters (Figure. 5B). Although the abundance of individual TFBS categories differed among genes, several transcription factor families were broadly represented throughout the gene family, suggesting that SlEPF/EPFL transcription may be controlled by a complex regulatory network involving multiple classes of transcription factors. Promoter-level comparisons further revealed considerable variation in the overall number of predicted motif-position associations. Among the analyzed genes, *SlEPF11* exhibited the highest number of predicted TFBS associations, whereas *SlEPF12* showed the lowest. Taken together, the pronounced variation in both cis-regulatory elements and predicted transcription factor-binding profiles suggests gene-specific regulatory characteristics within the *SlEPF* family, potentially enabling differential transcriptional responses to developmental, hormonal, and environmental cues.

### Tissue-specific expression profiles of *SlEPF* genes

Tissue-specific expression profiles of the *SlEPF* gene family were examined using publicly available RNA-seq data obtained from the BAR database at the University of Toronto (Figure 6) [57]. Expression patterns varied markedly among family members, indicating substantial tissue-dependent regulation. *SlEPF6* showed the highest expression level among the analyzed genes, with particularly strong accumulation in leaves (RPKM=43.07), together with detectable expression in flower buds and flowers. Similarly, *SlEPF11* exhibited high transcript abundance in leaves (RPKM=34.75) and relatively elevated expression in flower buds and flowers. *SlEPF7* and *SlEPF10* were preferentially expressed in reproductive tissues, with pronounced transcript accumulation in flower buds and flowers. In particular, *SlEPF7* reached RPKM values of 20.51 and 16.03 in flower buds and flowers, respectively, whereas *SlEPF10* displayed expression levels of 13.93 and 16.15 in the same tissues.

**Figure 6.**
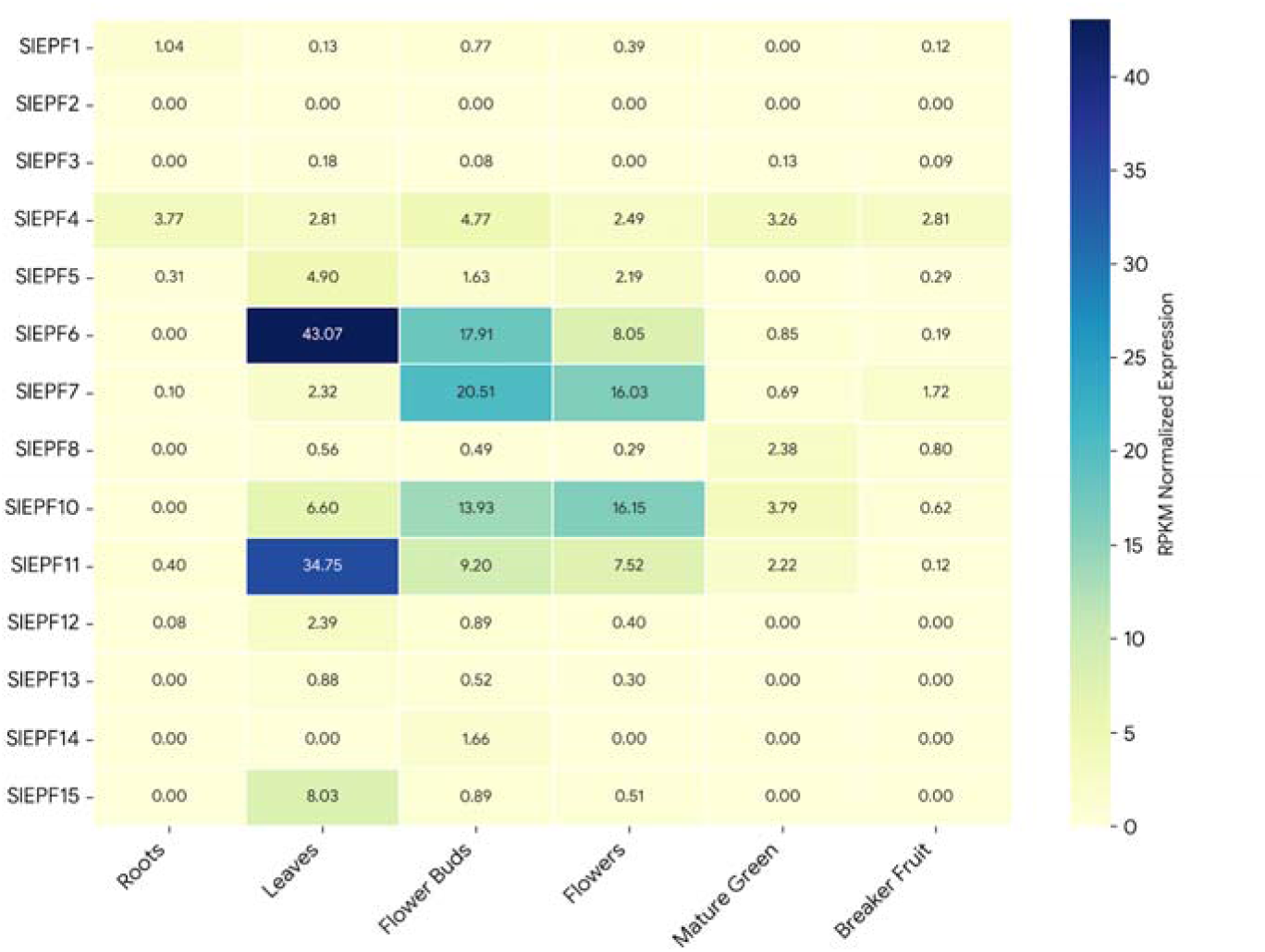
Tissue-specific expression profiles of *SlEPF* genes in *Solanum pimpinellifolium*. Transcript abundance of *SlEPF* family members across roots, leaves, flower buds, flowers, mature green fruit, and breaker-stage fruit was visualized as a heatmap using publicly available RNA-seq data obtained from the BAR database. Expression values are presented as RPKM-normalized transcript levels, with color intensity indicating relative expression abundance. Numerical values within each cell represent the corresponding RPKM values for each gene-tissue combination. Expression data for *SlEPF9* were not available in the database and were therefore excluded from the analysis.

Several other family members exhibited more restricted or comparatively low expression profiles. *SlEPF15* showed preferential expression in leaves, whereas *SlEPF5* was also most abundant in leaf tissue. In contrast, *SlEPF4* displayed relatively consistent expressions across all examined tissues, suggesting a broader expression pattern. Transcript levels of *SlEPF2*, *SlEPF12*, *SlEPF13*, and *SlEPF14* were generally very low or undetectable in most tissues included in the dataset. Overall, expression levels were generally lower in fruit tissues, particularly at the breaker stage, compared with vegetative and floral tissues. Expression data for *SlEPF9* were not available in the BAR dataset and were therefore excluded from the analysis.

### Expression responses of *SlEPF* genes to drought and high-temperature stress

Investigating the expression patterns of selected *EPF* genes in two tomato genotypes Money Maker (MM) and Ailsa Craig (AC) under different abiotic stress conditions provides valuable insights into their potential roles in plant stress responses. The expression analysis revealed distinct transcriptional responses of five putative abiotic stress-responsive genes (*SlEPF5, SlEPF8, SlEPF9, SlEPF10,* and *SlEPF13*) in MM under drought, heat, and combined drought–heat stress (Figure 7). Under drought stress, *SlEPF5* and *SlEPF13* were strongly downregulated, with log□-fold changes of −4.6 and −4.4, respectively. In contrast, *SlEPF8* was markedly upregulated (2.9), whereas *SlEPF9* and *SlEPF10* exhibited mild downregulation of −1.3 and −0.3, respectively. Under heat stress, *SlEPF5*, *SlEPF9*, and *SlEPF13* were downregulated, with log□-fold changes of −0.3, −0.8, and −3.8, respectively. *SlEPF8* showed a negligible change in expression (0.1), while *SlEPF10* was slightly upregulated (0.5). Among these genes, *SlEPF13* exhibited the strongest repression under heat stress. Under combined drought–heat stress, *SlEPF5* and *SlEPF13* remained strongly downregulated, with log□-fold changes of −2.7 and −4.4, respectively. *SlEPF9* showed only a minor reduction (−0.1), whereas *SlEPF8* and *SlEPF10* were upregulated, with values of 2.5 and 0.7, respectively. Additionally, the expression profiles of the same genes were investigated in AC under the same abiotic stress conditions (Figure 7). The results revealed predominantly positive transcriptional responses, although expression patterns varied among genes and stress conditions. Under drought stress, *SlEPF5* and *SlEPF13* were downregulated, with log□-fold changes of −1.7 and −1.9, respectively. In contrast, *SlEPF8* exhibited strong upregulation (2.8), while *SlEPF9* and *SlEPF10* were moderately upregulated, with values of 0.8 and 1.2, respectively. Under heat stress, all five genes were upregulated. *SlEPF10* showed the strongest induction, with a log□-fold change of 4.3, followed by *SlEPF9* (2.9), *SlEPF8* (2.6), *SlEPF13* (2.2), and *SlEPF5* (1.9). These results indicate a broadly positive transcriptional response of the examined *SlEPF* genes to heat stress in AC. Under combined drought–heat stress, *SlEPF8* and *SlEPF10* exhibited the strongest induction, with log□-fold changes of 5.1 and 4.4, respectively. *SlEPF9* was also substantially upregulated (2.7), followed by *SlEPF5* (1.4). In contrast, *SlEPF13* showed only a minor change in expression (−0.2), indicating a nearly stable transcriptional profile under combined stress. These patterns suggest that *SlEPF*-associated transcriptional responses in AC are strongly dependent on the type of abiotic stress.

**Figure 7.**
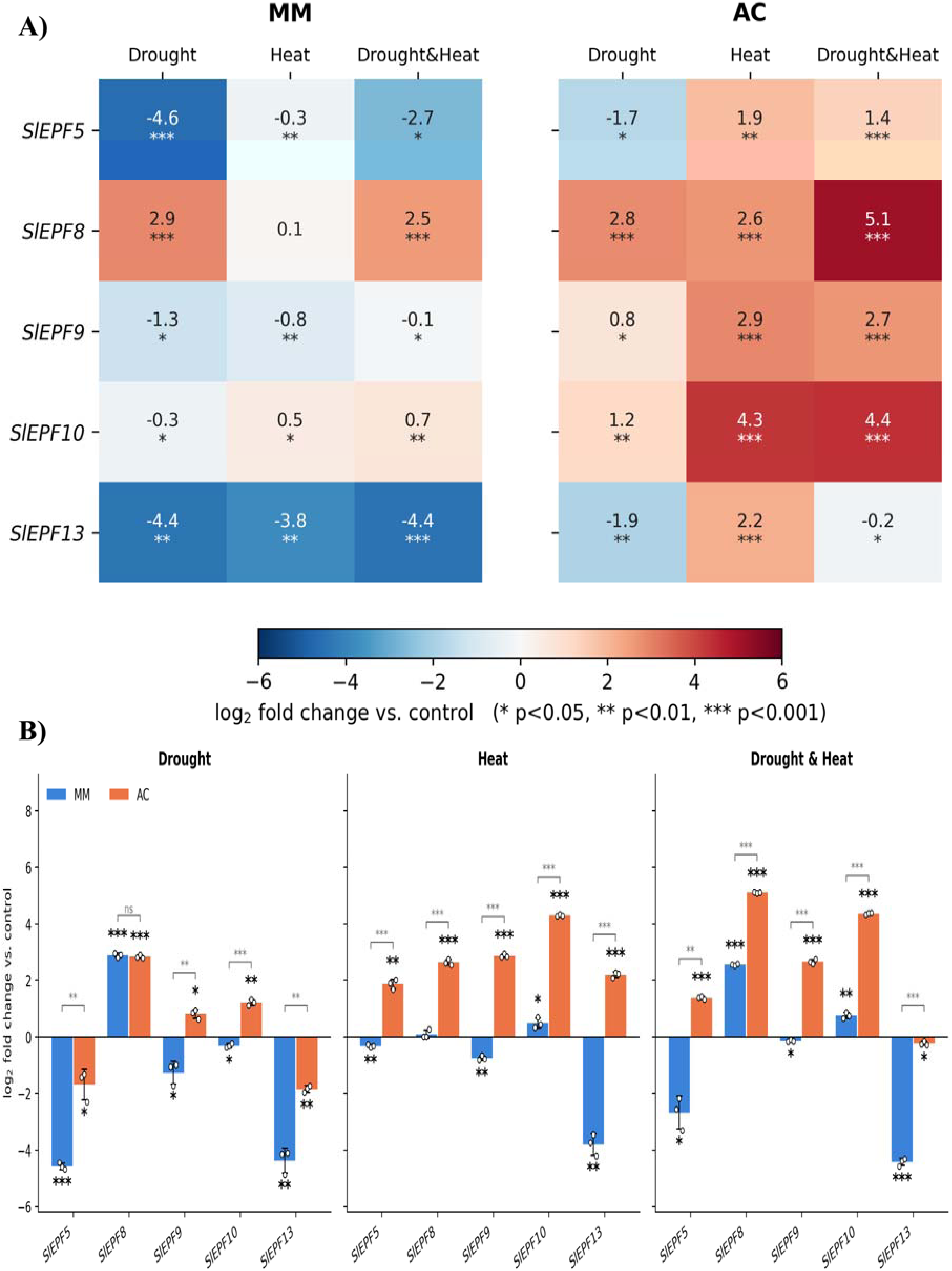
A) Heatmap of the relative expression of *SlEPF5*, *SlEPF8*, *SlEPF9*, *SlEPF10* and *SlEPF13* in genotypes Money Maker (MM) (left panel) and Ailsa Craig (AC) (right panel), determined by RT-qPCR. Within each panel, columns show the three stress treatments (drought, heat, and drought & heat) and rows show the genes. Expression values were calculated by the 2^-ΔΔCt method relative to the untreated control of the same genotype (control = 1) and are shown as log□-fold change, where 0 means no change. The number in each cell is the mean log□-fold change of three biological replicates. The color scale runs from blue (down-regulation) through light grey (no change) to red (up-regulation), with the same scale in both panels so the two genotypes can be compared directly. Asterisks below the numbers mark a significant difference from the control, tested with a one-sample *t*-test on log values against 0: * *P* < 0.05, ** *P* < 0.01, *** *P* < 0.001. Cells without asterisks did not differ significantly from the control. B) Bars show the mean of three biological replicates (n = 3), error bars show the standard deviation (SD) and open circles show the individual replicates. Asterisks above the bars indicate a significant difference from the control (one-sample *t*-test of log2 fold change values against 0). Grey brackets indicate a significant difference between MM and AC for the same gene and condition (Welch’s *t*-test). * *p* < 0.05, ** *p* < 0.01, *** *p* < 0.001, ns = not significant. The figure was generated in Python.

## Discussion

The effects of global warming, largely driven by rising atmospheric CO□ concentrations, have become increasingly pronounced, with significant implications for agricultural productivity and ecosystem stability [60]. At first glance, elevated CO□ concentrations may appear beneficial to plants because CO□ is an essential substrate for photosynthesis, and its increased availability can enhance photosynthetic rates and plant growth under certain conditions. However, this response is not universal, as the benefits of elevated CO□ are constrained by factors such as nutrient availability, water supply, temperature, and plant physiology [61, 62]. Under these conditions, plants exhibit developmental plasticity that enables them to adjust photosynthetic activity (CO□ assimilation) and water loss through the modulation of leaf traits as new leaves emerge. Stomatal ontogenesis, patterning, and function are essential components of plant adaptation to environmental stresses. The establishment and spatial distribution of the stomatal cell lineage, followed by its terminal differentiation into guard cells, are regulated by a key protein phosphorylation cascade involving the YODA mitogen-activated protein kinase kinase kinase (MAPKKK) [63-65]. In *Arabidopsis thaliana*, a well-characterized genetic regulatory network governs stomatal development and patterning. This network involves several basic *helix–loop–helix (bHLH)* transcription factors, including *SPEECHLESS (SPCH)*, *MUTE, FAMA*, and the *ICE/SCREAM (ICE/SCRM)* family, which regulate successive stages of stomatal lineage progression and differentiation [31, 66-69]. Their activities are coordinated through signaling pathways involving mobile *EPIDERMAL PATTERNING FACTOR (EPF)* peptides, which modulate stomatal development and ensure appropriate spacing between stomata [70]. Although *EPF* family members play an irreplaceable role in regulating stomatal development and may contribute to plant responses to abiotic stress, their functions in *S. lycopersicum*, particularly under stress conditions, remain poorly characterized.

In this study, we identified 15 members of the *EPF* gene family in the tomato genome. This number is slightly higher or lower than the number of *EPF* family members identified in other plant species, such as cotton species; *G. raimondii* (20) and *G. arboreum* (24) [71], wheat species (365) [41], *Populus trichocarpa* (15) [38], *Solanum tuberosum* L. (14) [37], *Secale cereale* L. (12) [72], *Sorghum bicolor* L. (12) [36], *Avena sativa* (33) [40], *Brassica napus* L. (27) [39], *Populus euphratica* (14) [73], *Nicotiana tabacum* (23) [74], *Brassica rapa* (14) [34], and *Zea mays* L. (16) [35]. The EPF domain was identified in all SlEPF proteins, except SlEPF6, which contains an EPF superfamily domain, and SlEPF11, which harbors a STOMAGEN-like family domain. In the remaining SlEPF proteins, the EPF domain was consistently located in the C-terminal region. Notably, 13 of the 15 SlEPF proteins possess a terminal proline residue, which may confer resistance to carboxypeptidase-mediated degradation and thereby contribute to peptide stability and the maintenance of a receptor-binding conformation. This result was aligned with Harald (2004) who illustrated that carboxypeptidase A preferentially cleaves C-terminal aromatic and aliphatic amino acid residues with long side chains, whereas glycine and acidic residues are released more slowly. In contrast, C-terminal lysine, arginine, proline, and histidine residues inhibit further cleavage by this enzyme [75]. Additionally, proline residues within C-terminal tails can restrict conformational mobility and contribute to correct folding, as shown for the potato carboxypeptidase inhibitor [76]. Most SlEPF proteins were predicted to contain three disulfide bonds, whereas SlEPF5 and SlEPF15 were predicted to contain two and four, respectively. The overall conservation of the cysteine pattern suggests that SlEPFs share a disulfide-stabilized scaffold, which is likely important for ligand activity. Ohki et al (2011) illustrated that STOMAGEN consists of a loop and a scaffold stabilized by three disulfide bonds. Swapping the loop and scaffold regions between STOMAGEN and EPF2 reverses their effects on stomatal density in *Arabidopsis* cotyledons, while deletion of a disulfide bond disrupts protein folding and abolishes activity [33].

Phylogenetic analysis of the *EPF* gene family across different plant species revealed two major clades (Clades A and B), which were further subdivided into distinct subgroups according to their evolutionary relationships. Clade A comprised six subgroups, whereas Clade B contained four (Figure 1B). AtEPF1 and AtEPF2 negatively regulate stomatal development, whereas AtEPFL9/STOMAGEN promotes stomatal formation, with these factors acting antagonistically to control stomatal density in leaves [77]. However, previous studies have reported a positive association between *EPF* gene expression and water-use efficiency (WUE), as demonstrated by the heterologous expression of the apple-derived *MdEPFL2* gene in tomato [78]. Therefore, the phylogenetic clustering of *SlEPF* members provides insights into their potential functional similarities within the same subgroup and establishes a basis for future investigations into the molecular mechanisms by which *EPF* genes regulate stomatal development in leaves.

The PPI network analysis revealed potential interactions between SlEPF proteins and several signaling components, including A0A3Q7G285 and A0A3Q7HPD8 (Serine/Threonine protein kinases) and A0A3Q7J9L5 (ERECTA leucine-rich repeat receptor-like Serine/Threonine protein kinase). For SlEPF13, the network also included A0A3Q7G821, an epidermal patterning factor-like protein. Notably, A0A3Q7G285 occupied a central hub position, suggesting a potential role in coordinating protein–protein interactions within the network. A second cluster comprised three interconnected proteins (A0A3Q7IQ87, A0A3Q7G3I6, and A0A3Q7GQI0), including a mitogen-activated protein kinase kinase kinase (MAPKKK), pointing to a possible link between EPF-associated signaling and MAPK-mediated regulatory pathways. Collectively, these predicted interactions suggest potential connections between EPF signaling, receptor-mediated perception, and downstream signaling cascades involved in stomatal development and stress responses; however, experimental validation is required to confirm these interactions and their functional significance. GO annotation analysis revealed that most of the identified genes were associated with biological processes related to stomatal complex development and guard cell differentiation. *SlEPF7*, *SlEPF13*, and *SlEPF15* were associated with the negative regulation of stomatal complex development. These findings are consistent with those of He et al. (2025), who reported that virus-induced gene silencing (VIGS) of *GhEPFL1-1*, *GhER1*, and *GhSERK17* resulted in increased stomatal density [79]. Additionally, He et al. (2025) reported that VIGS of *GhEPF2–2* increased stomatal density, whereas silencing of *GhEPFL9–6* resulted in a reduction in stomatal density, highlighting the contrasting roles of these genes in regulating stomatal development [80]. Moreover, in rice, overexpression of *OsEPF1* or *OsEPF2* substantially reduced stomatal density, while ectopic expression of these genes in *Arabidopsis* mutants (*epf2* and *epf1 epf2*) partially or significantly suppressed their elevated stomatal density. Conversely, silencing *OsEPFL9* in rice resulted in fewer stomata than in wild-type plants, whereas heterologous overexpression of *OsEPFL9* in *Arabidopsis* promoted excessive stomatal formation [81]. Regarding molecular functions, *SlEPF* genes were associated with MAP kinase activity, receptor serine/threonine kinase binding and receptor ligand activity. In particular, *SlEPF11* and *SlEPF13* were associated with protein kinase binding, whereas *SlEPF11* was additionally linked to receptor serine/threonine kinase binding. Furthermore, *SlEPF7*, *SlEPF8*, and *SlEPF12* were associated with receptor ligand activity. These findings are consistent with previous studies demonstrating that EPF peptides interact with receptor-like kinases, including receptor-like serine/threonine protein kinases, to regulate stomatal development [27, 80].

Signal peptide prediction identified an N-terminal secretion signal in 8 of the 15 SlEPF proteins with confident cleavage sites (SlEPF1, SlEPF2, SlEPF3, SlEPF8, SlEPF9, SlEPF10, SlEPF11 and SlEPF12). SlEPF15 was flagged with an uncertain call, and six proteins (SlEPF4, SlEPF5, SlEPF6, SlEPF7, SlEPF13 and SlEPF14) had no predicted signal peptide. The confident cleavage sites lie between residues 22-23 (SlEPF3) and 35-36 (SlEPF10), within the usual 20-35 residue range of plant secretory signal peptides. This agrees with the pre-pro-peptide architecture of EPF ligands in *Arabidopsis*, where the signal peptide directs the precursor to the secretory pathway and the cysteine-rich C-terminal domain is processed into the mature apoplastic peptide [77, 82]. It also agrees with the enrichment of extracellular region in the STRING analysis (panel C).

Promoter analysis of the 15 *SlEPF* genes identified a broad repertoire of CREs, which suggests that the tomato *EPF* family is regulated at the transcriptional level by developmental, environmental and hormonal cues. Beyond the core promoter elements (TATA-box and CAAT-box), the remaining CREs fell into three functional groups: growth and development (177 elements), stress responsiveness (166) and phytohormone responsiveness (112). Every *SlEPF* promoter carried elements from all three groups, but their composition differed markedly between genes. This indicates functional diversification within the family, consistent with the distinct, and in part antagonistic, roles of EPF peptides in stomatal patterning. The predominant expression of most *SlEPF* genes in leaf tissues is consistent with our hypothesis regarding their potential roles in regulating stomatal development and function. Moreover, this observation is consistent with the established role of the *CHALLAH* subfamily in promoting plant growth and development [83, 84]. Additionally, these findings are consistent with previous studies in *Sorghum bicolor*, Potato, *Gossypium*, and *Populus euphratica* demonstrating that the promoter regions of these genes frequently harbor cis-regulatory elements associated with abiotic stress responses and phytohormone signaling, highlighting their potential roles in coordinating plant development and stress adaptation [36, 37, 71, 73].

The expression profiles of the five selected *SlEPF* genes revealed distinct responses to different stress conditions. Under drought stress, three genes *(SlEPF8, SlEPF9,* and *SlEPF10*) were upregulated in the drought-tolerant AC cultivar, suggesting their potential involvement in drought stress responses. Similarly, Liu et al. (2024) reported that most *PtEPF* family members were markedly upregulated under drought stress, suggesting their potential roles in drought-responsive mechanisms [38]. Another study provided detailed insights into the *StEPF* gene family, demonstrating that *StEPF4* plays a crucial role in regulating drought resistance in potato by reducing stomatal density [37]. Under heat and combined drought–heat stress, all five *SlEPF* genes were upregulated, except for *SlEPF13*, which was downregulated under combined drought–heat stress. These findings are consistent with Negi et al. (2025), who demonstrated that the coordinated but contrasting expression of the antagonistic regulators *SoEPF9* (upregulated) and *SoEPF2* (downregulated) contributed to the open, small, and dense stomatal phenotype in *Sugarcane* mutant M4209, resulting in enhanced stomatal conductance, transpirational cooling, and gaseous exchange [85]. Furthermore, genetic analyses have demonstrated an epistatic relationship between YODA and HSP90.1, suggesting that these proteins act sequentially within the same developmental pathway to regulate stomatal formation. HSP90 proteins interact with YODA, influence its cellular polarization, and modulate the phosphorylation of downstream signaling components, including MPK6 and SPCH, under both normal and heat-stress conditions [65]. Beyond their roles in abiotic stress responses, *EPF* genes are also involved in plant growth and development, including root development and chloroplast-related processes [86, 87]. Additionally, Luling et al. (2022) reported that the loss of function of most *OsEPF* genes caused no obvious phenotypic alterations, whereas disruption of *OsEPFL2* in rice cultivar Kasalath resulted in shortened or absent awns and reduced grain size. The strong expression of *OsEPFL2* in young panicles further supports its potential role in awn development and grain formation [88]. Beyond their roles in abiotic stress responses, *EPF* genes also contribute to plant responses to biotic stress. Previous studies have reported differential expression of *EPF* family members in response to phytopathogenic fungal infections across various plant species. For example, *Botrytis cinerea* infection significantly downregulated the expression of six genes encoding predicted EPFL peptides in the moss *Physcomitrium patens*, while the expression of other family members remained unchanged [89]. In tomato, biotic stress differentially modulates *SlEPF* gene expression. Infection with *Fusarium oxysporum f. sp. lycopersici* upregulates *SlEPF7* while downregulating *SlEPF1* and *SlEPF5*. Similarly, elicitors from the non-pathogenic strain *Fusarium sambucinum* induce *SlEPF6/7* expression but suppress *SlEPF3/5* [90]. These diverse functions highlight the biological importance of the *EPF* gene family and underscore the need for further functional characterization to elucidate their regulatory mechanisms and potential applications in developing climate-resilient crops and promoting sustainable agriculture.

## Conclusion

In conclusion, the comprehensive characterization of the *SlEPF* gene family identified 15 members in tomato (*S. lycopersicum* L.), revealing conserved structural features and evolutionary relationships associated with stomatal development. Functional annotation and promoter analysis further highlighted their potential involvement in stomatal regulation, plant development, and environmental stress responses. Moreover, the contrasting expression profiles of five selected genes (S*lEPF5, SlEPF8, SlEPF9, SlEPF10*, and *SlEPF13*) between the heat-tolerant drought-sensitive Money Maker (MM) and semi-heat-tolerant Ailsa Craig (AC) tomato genotypes demonstrated genotype- and stress-dependent transcriptional regulation. Notably, the additive induction of *SlEPF8* in AC under combined stress conditions suggests its potential contribution to abiotic stress responses. These findings provide a foundation for further functional characterization of *SlEPF* genes and elucidation of their roles in stomatal regulation and tomato adaptation to drought, heat, and combined stress conditions.

## Data Availability Statement

All data generated or analyzed during this study are included in the article and its supplementary materials. Further inquiries can be directed to the corresponding author.

## Acknowledgments

The authors thank the support of the COST Action RECROP CA22157 (COST-European Cooperation in Science and Technology) for networking activities and Short-Term Scientific Mission (STSM). We acknowledge the Core Facility Plant Sciences of CEITEC Masaryk University for their support in obtaining scientific data presented in this paper.

## Ethics Approval

Not applicable.

