## Supplemental Table 1 for "EPF/EPFL Peptide Signaling as a Regulatory Node Linking Stomatal Development to Drought and Heat Tolerance in Tomato (*Solanum lycopersicum* L.)": Supplementary file 2.pdf

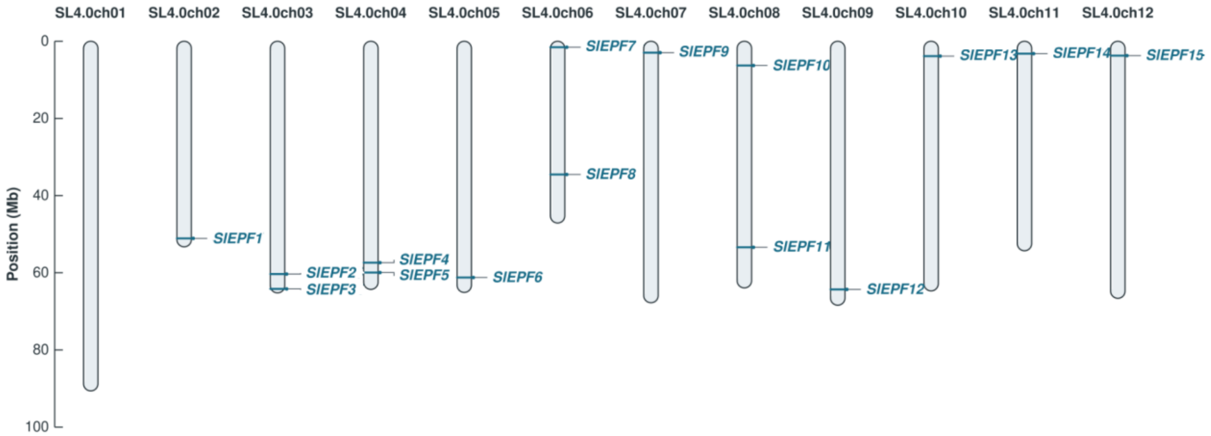


**Figure S1. Chromosomal distribution of the *SlEPF* genes in tomato.** The 15 *SlEPF* genes are mapped onto the 12 chromosomes of the tomato genome (SL4.0 assembly). Chromosome numbers are shown above each bar, and the scale on the left indicates physical position in megabases (Mb). Horizontal lines mark the approximate gene positions, and gene names are given beside them.
